# Mural β3-integrin signposts Stroma6, a gene signature predicting early relapse in low-risk and chemoresistant breast cancer

**DOI:** 10.64898/2026.09.10.750700

**Authors:** Alexander M Jordan, Vimala Anthonydhason, Katie Baird, Eleni Maniati, J. Louise Jones, Isobelle Wall, Anita Grigoriadis, Jun Wang, Tarek Abdel-Fatah, Kairbaan Hodivala-Dilke

## Abstract

The Oncotype DX (ODX) assay guides adjuvant chemotherapy in ER^+^/HER2^−^ breast cancer but overlooks drivers from the tumour stroma, leaving some low-risk and chemotherapy-spared patients vulnerable to rapid relapse. Here, we show that perivascular mural β3-integrin protein expression predicts early (3-year) relapse in ODX low-risk patients (HR = 15.80, *P* = 0.010), from which we derived a translatable 6-gene stromal signature, termed Stroma6. High Stroma6 expression predicted early recurrence in the ODX low-risk discovery cohort, with zero early relapses in the low-expression group. External validation in the METABRIC dataset confirmed that Stroma6 independently predicts rapid recurrence in ODX low-risk patients (HR = 11.68, *P* = 0.002). Furthermore, Stroma6 utility extends to clinical chemoresistance, independently predicting early events in a pan-subtype cohort with residual disease following neoadjuvant chemotherapy (HR = 2.29, *P* = 0.003). Ultimately, Stroma6 is a robust prognostic signature identifying patients at high risk of early recurrence across diverse clinical contexts.

## Introduction

Optimising adjuvant chemotherapy usage in hormone receptor-positive (HR^+^), human epidermal growth factor receptor 2-negative (HER2^−^) breast cancer (BC) requires precise stratification to prevent both overtreatment and undertreatment. This decision is largely guided by genomic assays, particularly Oncotype DX (ODX)^1^. While landmark trials, including TAILORx^2^ and RxPONDER^3^, established that many patients with low to intermediate Recurrence Scores (RS) can safely avoid adjuvant chemotherapy, a clinically significant minority of these patients still experience rapid relapse, representing a major challenge in precision oncology^4,5^. Critically, since the ODX assay profiles tumour cell intrinsic genes^6^, it inherently overlooks prognostic drivers arising from the tumour stroma.

The vascular niche forms a key part of the tumour stroma, where blood vessels (BVs) can promote tumour growth, metastasis and therapy resistance through angiocrine (endothelial cell) and pericrine (mural cell) signalling^7–10^. We have previously demonstrated that perivascular mural cell beta3-integrin (β3-integrin) protein expression regulates tumour growth in late-stage ER^+^ BC through altered pericrine signalling^11^. However, the influence of mural β3-integrin on local stromal transcriptomic profiles and its association with early recurrence in early-stage ER^+^/HER2^-^ BC remains unexplored. Here, we identify that mural β3-integrin protein expression signposts a high-risk tumour stroma-derived 6-gene signature, Stroma6, that predicts early relapse in patients classified as low-risk by ODX and validates in the chemoresistant neoadjuvant setting.

## Results

### Mural β3-integrin sub-stratifies prognosis in ODX low-risk breast cancer

To evaluate the prognostic value of perivascular mural β3-integrin, we utilised tissue microarrays (TMAs) from the ODX Nottingham University Hospital (NUH) discovery cohort, comprising early-stage, neoadjuvant treatment-naïve ER^+^/HER2^−^ BC patients with known ODX RS (***Figure 1a*; *Extended Data Table 1***). Cell DIVE multiplex immunofluorescence (MxIF) was performed to detect β3-integrin, CD31 (endothelial cells) and αSMA (mural cells). Using an AI-based classifier (HALO AI; F1 score = 0.91; ***Extended Data Table 2***), β3-integrin protein intensity was directly quantified within αSMA^+^ mural cells adjacent to CD31^+^ BVs (***Extended Data Figure 1a***). Patients were stratified into mural β3-integrin ‘high’ and ‘low’ expression groups using maximally selected log rank statistics. To define the clinical context of this biomarker, relapse-free survival (RFS) was assessed within distinct ODX risk categories (low, intermediate, high) and by adjuvant chemotherapy status, as this captures the real-world clinical consensus on patient risk.

**Figure 1.**
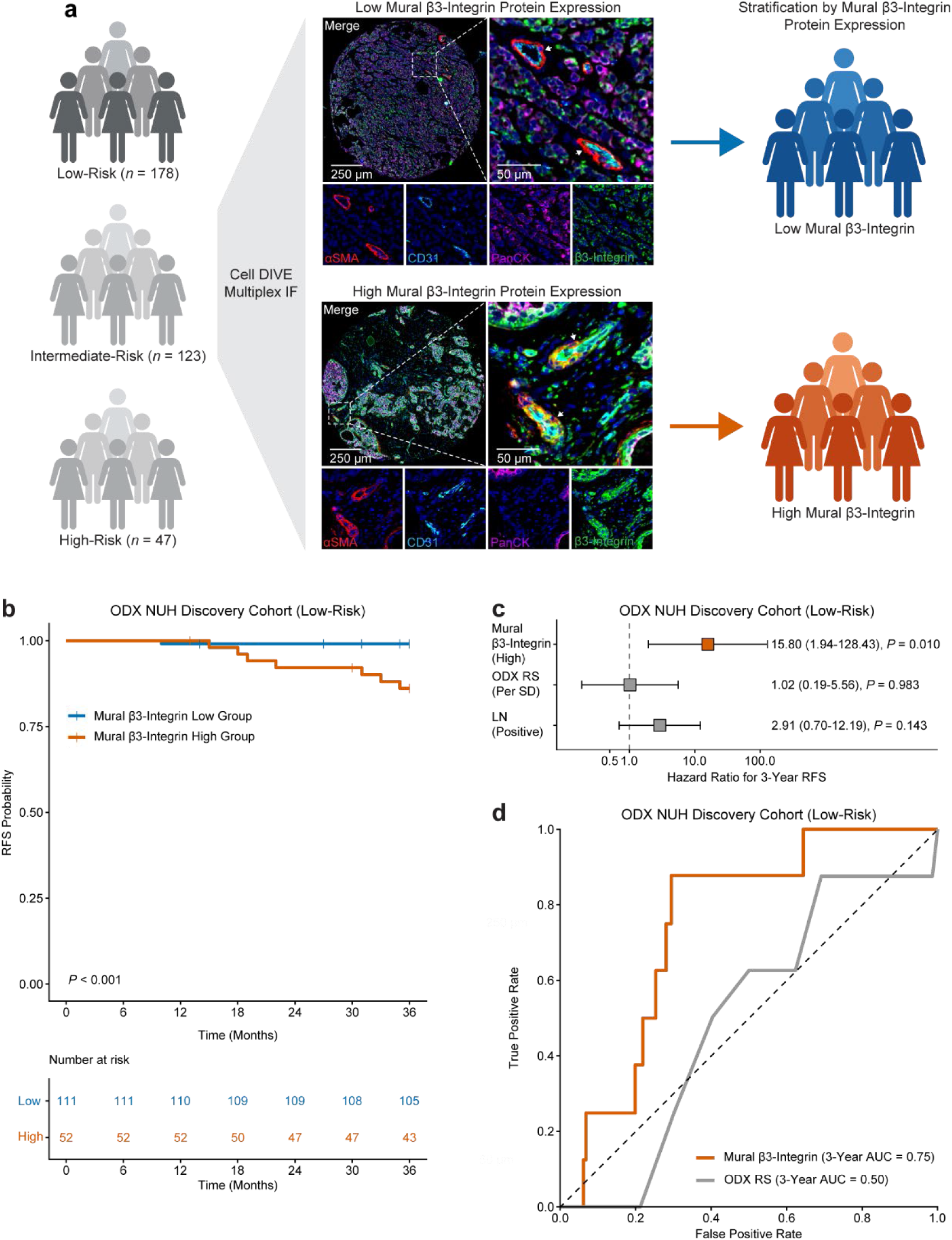
High mural β3-integrin protein expression predicts early relapse in Oncotype DX low-risk breast cancer. **a**, Schematic workflow and representative Cell DIVE multiplex immunofluorescence (MxIF) of the Oncotype DX (ODX) Nottingham University Hospital (NUH) discovery cohort (*n* = 348). The cohort comprised three clinical ODX risk groups (low, intermediate, and high), which underwent spatial profiling and were subsequently stratified into low and high mural β3-integrin protein expression groups. Representative MxIF images (scale bars = 250 µm; insets = 50 µm) show αSMA^+^ (red) mural cells, CD31^+^ (aqua) endothelial cells, PanCK^+^ (magenta) tumour cells, and β3-integrin (green). αSMA^+^ tumour blood vessels (BVs) are indicated by white arrows. **b**, Kaplan–Meier 3-year relapse-free survival (RFS) for high (orange) and low (blue) mural β3-integrin in the ODX low-risk cohort (*n* = 163). *P* value by two-sided log-rank test. **c**, Univariable Cox regression for 3-year RFS (*n* = 163). HR shown for high mural β3-integrin (reference: low), positive lymph node (LN) invasion (reference: negative), and ODX recurrence score (RS), where HR reflects a 1-standard deviation (SD, 11.1 points) increase. Error bars represent 95% confidence intervals. **d**, Time-dependent receiver operating characteristic (ROC) curves comparing continuous mural β3-integrin (orange) and ODX RS (grey) for 3-year relapse (*n* = 163). AUC, area under the curve. Patients were stratified into high and low mural β3-integrin groups using cut-off points derived by maximally selected rank statistics.

Survival analysis revealed that high mural β3-integrin protein expression was associated with an increased risk of early (3-year) relapse specifically within the ODX low-risk cohort (*n* = 163, *P* < 0.001; ***Figure 1b***). Similarly, high mural β3-integrin predicted early relapse in all ODX discovery cohort patients managed exclusively without adjuvant chemotherapy (*n* = 232, 3-year RFS, *P* = 0.002; ***Extended Data Figure 2a***). Conversely, mural β3-integrin lacked prognostic utility in ODX-stratified high-risk (*n* = 42, 3-year RFS, *P* = 0.131) or chemotherapy-treated patients (*n* = 70, 3-year RFS, *P* = 0.163; ***Extended Data Table 3***). Evaluation of ODX intermediate-risk patients was not possible due to event sparsity (*n* = 3 events for 97 patients).

Univariable Cox regression confirmed high mural β3-integrin as a strong predictor of 3-year relapse in both the ODX low-risk patients (*n* = 163, 3-year Hazard Ratio (HR) = 15.80, 95% confidence interval (CI): 1.94–128.43, *P* = 0.010, ***Figure 1c***) and in patients who did not receive adjuvant chemotherapy (*n* = 232, 3-year HR = 7.59, 95% CI: 1.61–35.75, *P* = 0.010, ***Extended Data Figure 2b***). Crucially, standard clinical metrics, including ODX RS and lymph node (LN) status, failed to predict early relapse in these same cohorts, indicating that mural β3-integrin outperforms these features (***Figure 1c* *and Extended Data Figure 2b***).

Time-dependent receiver operating characteristic (ROC) analysis was performed to confirm the prognostic weight of mural β3-integrin as a continuous biomarker. Within the ODX low-risk cohort, mural β3-integrin intensity exhibited strong discriminatory accuracy for early relapse (*n* = 163, 3-year area under the curve (AUC) = 0.75, ***Figure 1d***). The predictive performance of high mural β3-integrin expression was mirrored in patients who received no adjuvant chemotherapy (*n* = 232, 3-year AUC = 0.70, ***Extended Data Figure 2c***). Given that the majority of these patients received standard adjuvant endocrine therapy (*n* = 296 of 302), these data indicate that high mural β3-integrin identifies an aggressive tumour phenotype primed for early clinical escape despite standard of care treatment.

### Mural β3-integrin protein expression signposts a tumour stroma-derived 6-gene signature that sub-stratifies prognosis in ODX low risk breast cancer patients

While high mural β3-integrin effectively identifies early-relapse patients within ODX low-risk and no adjuvant chemotherapy subsets, routine clinical application of spatial MxIF is unlikely. Therefore, translation of this mural β3-integrin protein phenotype into a clinically accessible gene signature was sought.

Using spatial transcriptomics (NanoString GeoMx) in ODX low-risk patient TMAs from the ODX NUH discovery cohort (***Figure 2a***), we identified epithelial cell (PanCK^+^), immune cell (CD45^+^) and stromal cell (αSMA^+^) compartment-specific differentially expressed genes (DEGs) between mural β3-integrin-high and -low tumours (***Extended Data Figure 3a–c***). Strikingly, only signatures derived from the stromal (αSMA^+^) compartment successfully stratified patient risk (***Extended Data Tables 4– 6***), reinforcing a stromal-driven relapse phenotype. Consequently, we focused on the stromal compartment, distilling its DEGs into an optimised 6-gene signature, that we have termed Stroma6 (continuous HR = 3.53, *P* = 0.023). Crucially, Stroma6 remained highly enriched in β3-integrin-high patients (***Figure 2b***), confirming successful translation of the underlying mural β3-integrin stratification.

**Figure 2.**
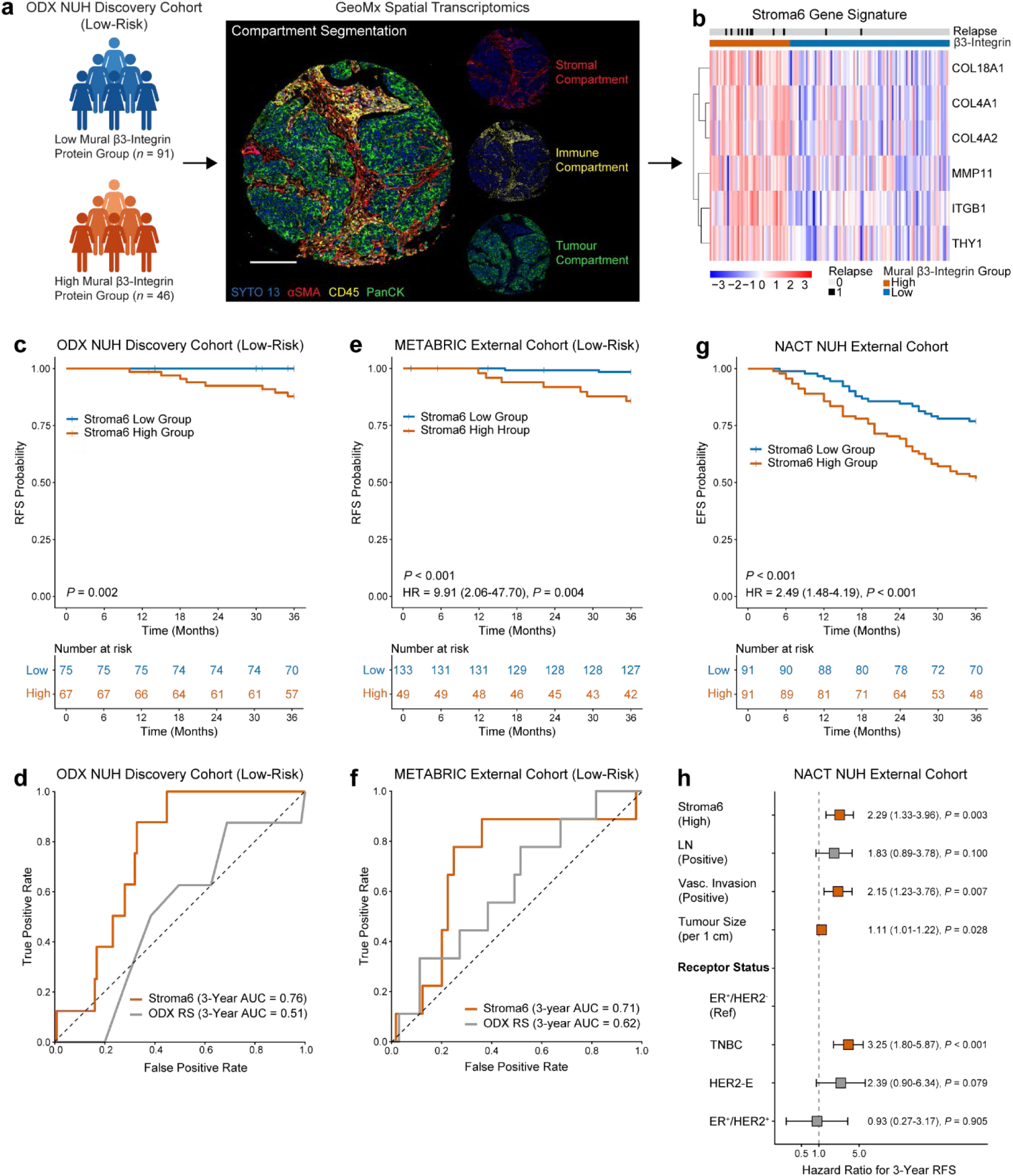
Stroma6 gene signature predicts early relapse in clinically low-risk and chemoresistant breast cancer. **a**, GeoMx spatial transcriptomics workflow for Stroma6 signature development. **b**, Heatmap of normalised Stroma6, 6-gene stromal signature expression (z-score) in the Oncotype DX (ODX) Nottingham University Hospital (NUH) discovery cohort low-risk group (*n* = 137), clustered by mural β3-integrin status (orange = high; blue = low). **c**, Kaplan–Meier (KM) 3-year relapse-free survival (RFS) for high (orange) and low (blue) Stroma6 in the ODX NUH discovery cohort low-risk group (*n* = 142). **d**, Time-dependent receiver operating characteristic (ROC) curves comparing continuous Stroma6 score (orange) and ODX recurrence score (RS; grey) for 3-year relapse in the ODX NUH low-risk discovery cohort (*n* = 142). AUC, area under the curve. **e**, KM 3-year RFS for high (orange) and low (blue) Stroma6 in the METABRIC ODX low-risk cohort (*n* = 182). Univariable Cox regression hazard ratios (HR) for high Stroma6 (reference: low) are shown. **f**, Time-dependent ROC curves comparing continuous Stroma6 score (orange) and ODX RS (grey) for 3-year relapse in the METABRIC ODX low-risk cohort (*n* = 182). **g**, KM 3-year event-free survival (EFS) for high (orange) and low (blue) Stroma6 in the neoadjuvant chemotherapy (NACT) NUH cohort (*n* = 182). **h**, Multivariable Cox regression for 3-year EFS in the NACT NUH cohort (*n* = 180). HR shown for high Stroma6 (reference: low), positive lymph node (LN) invasion (reference: negative), positive vascular invasion (reference: negative), tumour size (per 1 cm), and receptor status (TNBC, HER2-enriched, ER^+^/HER2^+^; reference: ER^+^/HER2^-^). Error bars represent 95% confidence intervals. For **c**, **e** and **f**, *P* value by two-sided log-rank test. For each cohort, patients were stratified into high and low Stroma6 groups using the global median Stroma6 score of the full, unstratified dataset.

To further evaluate the prognostic value of Stroma6, we stratified the whole ODX NUH discovery cohort by the median Stroma6 signature score. High Stroma6 expression was associated with significantly worse RFS in both the ODX low-risk subgroup (*n* = 142, 3-year RFS, *P* = 0.002; ***Figure 2c***) and in patients spared adjuvant chemotherapy (*n* = 206, 3-year RFS, *P* = 0.006; ***Extended Data Figure 4a***). As Stroma6 achieved perfect stratification in the ODX low-risk cohort – with zero events in the low-expression group – standard categorical Cox modelling was precluded. However, for patients who did not receive adjuvant chemotherapy, univariable Cox regression confirmed high Stroma6 expression as a potent predictor of 3-year relapse (*n* = 206, HR = 10.38, 95% CI: 1.31–81.90, *P* = 0.030; ***Extended Data Figure 4b***). Furthermore, time-dependent ROC analysis highlighted the strong discriminatory accuracy of Stroma6 for identifying early relapse, achieving a 3-year AUC of 0.76 in the ODX low-risk group (*n* = 142; ***Figure 2d***) and 0.73 in the no adjuvant chemotherapy group (*n* = 206; ***Extended Data Figure 4c***). As expected, Stroma6 provided no prognostic value in ODX intermediate-risk, ODX high-risk, or chemotherapy treated patients (***Extended Data Table 7***).

### Validation of the Stroma6 gene signature in external cohorts

To validate these findings, we evaluated the Stroma6 signature in the METABRIC cohort^12^. As this dataset utilises bulk transcriptomic data, this analysis also served to test the feasibility of translating our spatial signature into a standard clinical genomic assay. Restricting the cohort to endocrine-treated patients meeting standard ODX testing criteria, we applied an in silico ODX classifier that successfully recapitulated clinical risk groupings (*n* = 784; ***Extended Data Figure 5a***).

Within the ODX low-risk subset of this curated METABRIC cohort, high expression of Stroma6 identified patients with significantly worse 3-year RFS (*n* = 182, *P* < 0.001; ***Figure 2e***). Similarly, high Stroma6 expression correlated with poor 3-year RFS in patients managed without adjuvant chemotherapy (*n* = 715, *P* = 0.006; ***Extended Data Figure 5b***). After adjusting for standard clinicopathological features, multivariable Cox regression confirmed Stroma6 as a significant, independent predictor of early relapse (3-year) within the ODX low-risk group (*n* = 182, HR = 11.68, 95% CI: 2.40–56.92, *P* = 0.002; ***Extended Data Figure 5c***) and the no adjuvant chemotherapy group (*n* = 682, HR = 1.85, 95% CI: 1.09–3.13, *P* = 0.022; ***Extended Data Figure 5d***).

Furthermore, time-dependent ROC analysis of the continuous Stroma6 score demonstrated strong discriminatory accuracy for 3-year relapse within the ODX low-risk subset (*n* = 182, AUC = 0.71; ***Figure 2f***), while also maintaining prognostic value in the broader cohort of patients not treated with adjuvant chemotherapy (*n* = 715, AUC = 0.62; ***Extended Data Figure 5e***). In line with results from the discovery cohort, Stroma6 was not prognostic in ODX intermediate-risk, ODX high-risk, or chemotherapy treated patients (***Extended Data Table 8***). Collectively, these data demonstrate that the Stroma6 gene signature functions effectively as a bulk genomic assay to identify high-risk patients currently misclassified as clinically low risk.

Having validated the clinical utility of Stroma6 in endocrine-treated, ER^+^/HER2^-^ BC, we next sought to determine if this aggressive stromal phenotype also confers prognostic risk in the context of clinical chemoresistance. Therefore, we evaluated the Stroma6 signature in a pan-subtype cohort of BC patients who failed to achieve pathological complete response (pCR) following NACT.

Applying the signature to spatial transcriptomic data derived strictly from the stromal (αSMA^+^) compartment, high Stroma6 expression stratified patients with significantly worse 3-year event-free survival (EFS) (*n* = 182, *P* < 0.001; ***Figure 2g***). After adjusting for standard clinicopathological features and clinical receptor status, multivariable Cox regression confirmed Stroma6 as a robust, independent predictor of events at 3 years (*n* = 180, HR = 2.29, 95% CI: 1.33–3.96, *P* = 0.003, ***Figure 2h***). Furthermore, time-dependent ROC analysis revealed that continuous Stroma6 score displayed a superior discriminatory performance compared with residual tumour size (3-year AUC = 0.65 vs 0.59, ***Extended Data Figure 6a***). Ultimately, these results expand the potential clinical utility of Stroma6, establishing it as a robust prognostic signature of aggressive, therapy-resistant disease.

## Discussion

Our findings underscore a critical limitation of ODX, where a reliance on tumour-intrinsic profiling inherently overlooks malignant drivers from the tumour stroma. Consequently, a subset of patients clinically stratified as low risk by ODX, and spared adjuvant chemotherapy based on real-world clinical consensus, still experience rapid relapse. We demonstrated that perivascular mural β3-integrin successfully stratifies these patients, identifying an aggressive stromal phenotype conferring high risk of early recurrence (≤ 3 years). This aligns with emerging evidence implicating vascular pericrine signalling as a regulator of tumour progression and therapy resistance^7,11^.

Since mural β3-integrin protein expression is unlikely to be a feasible test in the clinical environment, we distilled this phenotype into the 6-gene Stroma6 signature, which validated in bulk transcriptomics. This provides a complementary tool to ODX amenable to standard reverse transcription quantitative real-time PCR (RT-qPCR) workflows. Capturing tumour stromal-derived drivers offers a strategy to identify high-risk patients who are currently undertreated, while preserving the de-escalation benefits of ODX for the true low-risk majority.

Furthermore, the signature’s prognostic utility extends beyond ODX low-risk, chemotherapy-spared patients. In a pan-subtype cohort with residual disease following NACT, high Stroma6 expression independently predicted early recurrence. This indicates that the Stroma6 signature identifies which chemoresistant tumours are primed for aggressive progression irrespective of BC subtype or treatment history.

This work has limitations. While validated across independent external cohorts, the Stroma6 signature was validated using bulk and spatial transcriptomics. Clinical translation will require validating the signature using standard RT-qPCR assays on tumour samples.

Genomic assays have undoubtedly spared thousands of patients from unnecessary chemotherapy. However, the next frontier of precision oncology requires integrating the tumour stroma into risk stratification. Our mural β3-integrin-derived Stroma6 signature bridges this gap, offering a novel approach to ensure that low risk truly implies safe from recurrence.

## Acknowledgements

AMJ support came from CRUK program grant (CRUK DRCNPG-May21/100004). VA is supported by the NIHR Barts Biological Research Centre (NIHR203330). KB and EM is supported by the CRUK City of London Major Centre (CTRQQR-2021\100004). KH-D, JW and JLJ are HEFCE supported through Queen Mary University of London. Other support for the lab includes Barts Charity (MGU0601), Medical Research Council (MR/V009621/1), and Worldwide Cancer Research (19-0108). IW was funded by the Wellcome Leap ΔTissue program. AG is funded by Breast Cancer Now [KCL-BCN-Q3], the Medical Research Council (MRC) [MR/X012476/1], Cancer Research UK [CRUK/07/012]. We thank Dr Gabriela Veronica D’Amico Lago, Queen Mary University of London, Barts Cancer Institute for her support of the study.

## Author contributions

AMJ: Conceptualization, validation, investigation, visualization, methodology, writing– original draft, project administration, writing–review and editing.

VM, KB, EM and JW: formal analysis, methodology, writing–review and editing. IW and AG: GeoMx resources and technical training, writing–review and editing. JLJ: Conceptualization and funding acquisition, writing–review and editing.

TA and KHD: Conceptualization, supervision, funding acquisition, project administration, writing–review and editing.

## Methods

### Patient Cohorts and Study Design

#### Discovery Cohort (Oncotype DX)

The retrospective discovery cohort initially comprised 348 patients with early-stage (Stage I–IIIA), ER^+^/HER2^−^ breast cancer (BC). Following exclusions due to tissue damage during processing or insufficient tumour blood vessel (BV) counts for spatial analysis, 302 patients were included in the final analytical cohort (***Extended Data Table 1***). All patients underwent clinical Oncotype DX (ODX) genomic testing to calculate Recurrence Scores (RS). Clinical risk categories were defined according to institutional practice at the time of diagnosis: low-risk (RS < 18), intermediate-risk (RS 18–30), and high-risk (RS > 30), with risk groups used to guide adjuvant chemotherapy decisions. Standard practice dictated that low-risk patients were spared adjuvant chemotherapy, high-risk patients were recommended chemotherapy, and intermediate-risk decisions were guided by clinicopathological features (primarily menopausal and lymph node status). Nearly all patients received adjuvant endocrine therapy (***Extended Data Table 1***). Primary formalin-fixed paraffin-embedded (FFPE) tumour tissue was obtained from surgical resections prior to any adjuvant therapy. Tissue microarrays (TMAs) were constructed using 1.3 mm diameter cores (80 cores per TMA block) and stored at 4°C under desiccated conditions.

#### External Validation Cohort 1 (METABRIC)

To validate the prognostic utility of the stromal signature in a bulk transcriptomic setting, clinical metadata and normalised microarray gene expression profiles were obtained from the Molecular Taxonomy of Breast Cancer International Consortium (METABRIC) dataset^12^. To strictly align with the discovery cohort and standard clinical genomic testing guidelines, the dataset was filtered to include only patients with early-stage (Stage 0–III, or unknown stage) ER^+^/HER2^−^ disease, endocrine therapy received, and zero to three positive lymph nodes (LN), resulting in a final analytical cohort of 785 patients (***Extended Data Table 1***). To evaluate the signature within clinically relevant contexts, in silico ODX RS were calculated for all patients (detailed in the Bioinformatics section) to assign them into matching low-, intermediate-, and high-risk clinical categories.

#### External Validation Cohort 2 (Neoadjuvant Chemotherapy)

To assess the prognostic utility of the stromal signature in the context of clinical chemoresistance, a secondary cohort comprising 182 pan-subtype BC patients was utilised (***Extended Data Table 1***). All patients in this cohort were treated with standard of care (SOC) neoadjuvant chemotherapy (NACT) prior to surgery. Inclusion criteria strictly required patients to have residual invasive disease – defined as a failure to achieve a pathological complete response (pCR) – at the time of surgical resection. FFPE residual tumour tissue was collected at surgery and used for spatial transcriptomic profiling. TMAs were constructed using 1.3 mm diameter cores (80 cores per TMA block) and stored at 4°C under desiccated conditions.

#### Ethics and Consent

Sample collection and utilisation for the discovery and NACT cohorts complied with all relevant ethical regulations and were approved by the Research Ethics Committee (REC; reference: 19/EM/0301; IRAS Project ID: 266220). Patient sample identifiers were fully anonymised and tracked within a centralised clinical database. Informed consent was obtained, or if a waiver granted?

### Multiplex Immunofluorescence Staining and Imaging (Cell DIVE)

TMA blocks for the ODX and NACT cohorts were sectioned at 5 µm thickness, floated in a 37°C water bath, and adhered to SuperFrost Plus slides. To ensure distinct morphology, increase BV sampling, and account for spatial heterogeneity, two sections taken a minimum of 25 µm apart were assessed for each TMA.

Multiplex immunofluorescence (MxIF) was performed using the Cell DIVE™ platform (Leica Microsystems) as previously described^13^, with specific modifications. Briefly, FFPE TMA sections were baked overnight at 56°C (12–18 h), deparaffinised, rehydrated, and permeabilised, followed by a two-step antigen retrieval process (pH 6 and pH 9). Sections were blocked using 10% goat serum (Sigma-Aldrich, G9023) for 1 h at room temperature (RT), then stained with DAPI (10 μg/ml) for 15 min at RT. Antibody incubation was performed in two rounds. Round 1 consisted of the unconjugated primary antibody anti-β3-integrin (Clone ERP17507, Abcam, ab240214; 1:400) incubated for 1 h at RT. Round 2 consisted of the secondary fluorescent antibody Goat anti-Rabbit IgG Alexa Fluor 750 (ThermoFisher Scientific, A-21039; 1:400) multiplexed with three directly conjugated primary antibodies: anti-αSMA Alexa Fluor 488 (Clone 1A4, Abcam, ab184675; 1:100), anti-CD31 Alexa Fluor 555 (Clone 89C2, Cell Signalling Technology, #61255; 1:80), and anti-PanCK Alexa Fluor 647 (Clone AE-1/AE-3, Novus Biologicals, NBP2-33200AF647; 1:100). Following each staining round, sections were washed three times for 5 min in wash buffer (0.01% Tween-20 in PBS).

Imaging was conducted using the Cell DIVE. A whole-slide image at 10x was acquired for region-of-interest (ROI) selection, followed by multiplex biomarker imaging at 20x. To account for intrinsic tissue autofluorescence (AF), baseline images of the TMAs were acquired in all appropriate fluorescent channels prior to antibody incubation. Following staining, the acquisition software automatically registered the biomarker images to the baseline DAPI stain to ensure precise spatial alignment. Computational AF subtraction was then applied to remove background tissue signal, generating highly specific, signal-to-noise optimised images for downstream spatial phenotyping.

### Image Analysis and Spatial Phenotyping

Digitised MxIF images from the ODX and NACT cohorts were imported into HALO image analysis software (Indica Labs). TMA cores were computationally segmented and quality control was performed to exclude severely damaged or folded tissue regions from downstream analysis.

To identify tumour BVs across heterogeneous tissue architectures, a deep learning semantic segmentation classifier was trained using HALO AI (DenseNet V2 algorithm). The network was trained using a binary classification approach to distinguish αSMA^+^ BVs from all other background tissue components. The model was trained at a resolution of 0.75 µm/pixel using iteratively annotated regions across multiple TMA cores to capture maximum morphological variance. A minimum object area threshold of 150 µm^2^ was applied to eliminate staining artefacts. The accuracy of the AI classifier was evaluated on a test set of four structurally heterogeneous TMA cores (encompassing highly cellular tumour regions, dense stroma, and varied BV morphologies), none of which were used to train the model. Semantic segmentation validation against manual ground-truth annotations yielded an F1 score of 0.91 (***Extended Data Table 2***), confirming high spatial accuracy.

Following BV segmentation, the HALO Area Quantification FL module (version 2.3.4) was utilised to spatially resolve distinct vascular compartments within the identified BVs. Compartments were defined based on pre-optimised pixel intensity thresholds (αSMA threshold: 7,000; CD31 threshold: 2,500). To capture the continuous spectrum of protein expression and prevent manual gating bias, no minimum intensity threshold was applied to the β3-integrin channel. The perivascular mural compartment was spatially defined as αSMA^+^/CD31^−^.

BV data were exported, and individual cores were mapped to patient identifiers. To account for the dual-section sampling strategy, BV data from both TMA sections were pooled for each patient prior to processing in R (version 4.3.1). To ensure robust spatial quantification and eliminate false positive segmentation fragments, individual BVs lacking an αSMA or CD31 area, or possessing a mural area of < 10 µm², were excluded. Finally, to generate robust patient-level biological representation, patients with fewer than 10 quantifiable BVs across their respective cores were excluded from downstream analysis. For each remaining patient, the median mural β3-integrin intensity across all valid BVs was calculated to yield a single representative score.

### Spatial Transcriptomics (NanoString GeoMx)

Spatial transcriptomic profiling was performed using the GeoMx Digital Spatial Profiler (DSP; NanoString Technologies). FFPE TMA sections for the ODX and NACT cohorts were baked at 60°C for 2 h, deparaffinised, and rehydrated. Target retrieval was performed by briefly immersing sections in boiling DEPC-treated water (10 s), followed by boiling 1X Tris-EDTA (pH 9.0) for 20 min. Tissues were washed in PBS and subsequently digested with Proteinase K (0.1 μg/ml) for 15 min at 37°C, washed, and post-fixed. Sections were then hybridised overnight (12–18 h) at 37°C with the GeoMx Whole Transcriptome Atlas (WTA) RNA probe panel using a HybEZ II Hybridisation System (ACDBio).

Following stringent washing (37°C) to remove off-target probes, sections were blocked in Buffer W (NanoString) for 30 min at RT. To guide spatial profiling and compartment segmentation, sections were stained with morphology markers via a three-round incubation protocol. Round 1 consisted of the unconjugated primary antibody anti-CD45 (Clone 2B11 + PD7/26, Novus Biologicals, NBP2-34287, 1:40) incubated for 1 h at RT. Round 2 consisted of the secondary fluorescent antibody Goat anti-Mouse IgG Alexa Fluor 594 (ThermoFisher Scientific, A-11032; 1:400) incubated for 1 h at RT. Round 3 consisted of the nuclear counterstain SYTO 13 (ThermoFisher Scientific, S7575, 1:1000), and the directly conjugated primary antibodies anti-PanCK (Alexa Fluor 532, Clone AE-1/AE-3, Novus Biologicals, NBP2-33200AF532; 1:390) and anti-αSMA (Alexa Fluor 647, Clone SP171, Abcam, ab267537, 1:1000) incubated for 1 h at RT. Following each staining round, sections were washed twice for 5 min in 2X SCC. Slides were then loaded onto the GeoMx DSP.

TMA cores were selected for profiling based on prior MxIF analysis; to ensure spatial transcriptomic data were generated for patients with corresponding mural β3-integrin analysis, cores exhibiting a high density of blood vessels were prioritised for ROI placement. ROIs were set to the maximum field of view (660 × 785 µm) and placed over each selected core. Within each ROI, areas of illumination (AOIs) were computationally segmented into discrete tumour (PanCK^+^), stromal (αSMA^+^), and immune (CD45^+^) compartments based on morphology marker expression. To ensure compartment purity and prevent spatial cross-contamination, custom morphological processing parameters were applied to each segment. AOIs containing fewer than 25 nuclei were excluded to ensure sufficient oligonucleotide collection. UV-cleaved indexing oligos from each segmented compartment were collected into 96-well plates and stored at −20°C.

For library preparation, collection plates were dried at 65°C for 1 h and rehydrated with nuclease-free water. PCR amplification was performed using the GeoMx Master Mix and Seq Code primers (NanoString) to add unique dual-indexing adapters to each sample. Following AMPure XP bead clean-up (Beckman Coulter), library quality, quantity, and fragment size (∼170 bp) were verified using a TapeStation system (Agilent). Libraries were subsequently pooled and outsourced for next-generation sequencing (NGS; Azenta Life Sciences).

### GeoMx Data Processing, Quality Control and Batch Correction

Raw FASTQ files were converted to digital count conversion (DCC) files using the *GeoMx NGS Pipeline* (2.3.3.10). *GeoMxTools*^14^ (3.5.0) was used to read in the DCC files resulting in expression data for 18,815 probes and 1817 samples including 20 No Template Control (NTC) samples. Probe annotation was obtained from the Homo sapiens Whole Transcriptome Atlas v1.0 PKC file provided on the NanoString website. Twenty-one negative probes previously identified as poorly performing across multiple cohorts and tissue types (FFPE, FF, TMA) were removed, leaving 18,794 probes for downstream analysis.

The standR^15^ (1.6.0) package was used to perform gene and sample-level quality control (QC). NTC QC was performed by assessing the total NTC counts and total raw counts for all probes. Since all NTC counts for all probes were uniformly relatively low between 0-3 counts, the segments were retained for further QC. Segment level QC was performed by excluding segments that had: rweights below the outlier thresholds of the robust linear models of the number of nuclei and area of illumination (AOI) size, sequence saturation percentage < 50% and a geometric mean of the negative probes <= 1 or > 100. Following segment level QC and exclusion of negative probes, there was expression data for 18,676 genes for 1768 samples. Genes with counts below the mean of the geometric mean of the negative probes in over 90% samples were discarded, leaving 13,143 genes for downstream analysis.

375 samples from 147 patients with both low-risk ODX designation and associated Cell DIVE β3-integrin protein intensity data were selected for analysis. Batch correction was performed with the *standR* package via the RUV4 method, with the combined variable of segment (αSMA^+^,CD45^+^, PanCK^+^) and mural β3-integrin group (low, high) as the factor of interest. Optimal k (k=1) was selected based on the silhouette scores using *findBestK()*.

### Derivation of the Stromal Gene Signature

Differential gene expression analysis comparing low and high β3-integrin group was performed with the *edgeR (*4.4.2)^16^ package using R version 4.4.2^17^. Normalisation factors were computed using *calcNormFactors()* with the trimmed mean of M values (TMM) method^18^. A design matrix was constructed incorporating the composite group variable of segment and mural β3-integrin group along with RUV4 batch correction factors. Low-coverage genes were filtered using *filterByExpr().* Differential expression was assessed using the *limma-voom*^19^ pipeline, applying *voomLmFit(), contrasts.fit(), and eBayes()*. Models were blocked by unique patient ID to account for repeated measurements P values were adjusted for multiple testing using the Benjamini-Hochberg method.

A weighted differential expression score (|log2FC| × −log10(*P*adj)) was used to rank genes, and the top 20 differentially expressed genes associated with mural β3-integrin status were selected across stromal (αSMA^+^), tumour (PanCK^+^), and immune (CD45^+^) compartments. Selected genes were Z-score normalised and visualised as hierarchically clustered heatmaps grouped by mural β3-integrin status.

To define compartment-specific signatures, candidate gene sets for stromal (αSMA^+^), tumour (PanCK^+^), and immune (CD45^+^) compartments were generated by sequentially aggregating the top 5–20 ranked genes. For each signature, a continuous score was computed per patient as the mean normalised expression of the selected genes. Although gene selection was derived from the ODX low-risk subset, signature scores were calculated across the full pooled GeoMx spatial transcriptomic dataset, including the discovery ODX cohort, validation NACT cohort, and additional spatial samples, ensuring comparability across clinical cohorts and minimising cohort-specific normalisation bias. Patients were then stratified into high- and low-signature groups using the median score within the ODX discovery cohort.

To determine the optimal prognostic model, we systematically evaluated each candidate signature (top 5 through 20 genes across all three compartments) specifically within the ODX low-risk subset. Prognostic performance (3-year relapse) for each candidate signature was evaluated using Kaplan-Meier (KM) analysis (log-rank test), time-dependent receiver operating characteristic (ROC) analysis, and univariable Cox proportional hazards regression for both continuous and categorical signature scores. Final model selection was determined by maximising predictive accuracy via Harrell’s Concordance Index (C-index) and minimising model complexity via the Akaike Information Criterion (AIC). Based on these evaluation criteria, the optimal 6-gene αSMA model was selected as the final stromal gene signature for all subsequent validation analyses.

### METABRIC Processing and In Silico Oncotype DX Scoring

Clinical metadata and normalised microarray gene expression profiles for the METABRIC cohort were accessed via the cBioPortal for Cancer Genomics^12^.

To evaluate the stromal biomarker within ODX risk classes, in silico Oncotype DX recurrence scores were calculated for all patients using the *genefu* R package^20^, enabling stratification into low-, intermediate-, and high-risk categories.

For transcriptomic validation, a continuous stromal signature score was calculated for each patient as the mean expression of the six signature genes. Patients lacking relapse-free survival (RFS) data were first excluded. Patients were then stratified into high- and low-expression groups based on the median signature score calculated across this METABRIC cohort. This ensured standardisation across clinical contexts, thereby preventing subgroup-specific normalisation bias. Following stratification, the dataset was strictly filtered to match clinical ODX testing guidelines: ER^+^/HER2^−^ receptor status, early-stage disease (Stage 0–III, or unknown stage), and zero to three positive LN, yielding the final analytical cohort.

### Statistical and Survival Analysis

All statistical analyses were performed using R (version 4.3.1). The primary clinical endpoints were RFS for the ODX and METABRIC cohorts, and event-free survival (EFS) for the NACT cohort. To specifically evaluate early and medium-term recurrence dynamics, survival times were strictly right-censored at 36 months (3-year) and 60 months (5-year) for all analyses.

For mural β3-integrin stratification, maximally selected rank statistics (surv_cutpoint function, survminer package) were used to determine the optimal prognostic cut-off point for continuous mural β3-integrin intensity. For the gene signature analyses, patients were stratified into ‘high’ and ‘low’ expression groups using the median continuous signature score. To ensure robustness and prevent cohort-specific thresholding bias during validation, the global median derived from the entire respective dataset was consistently applied across all clinical subsets.

Survival probabilities were estimated using the KM method, and differences between groups were assessed using the two-sided log-rank test (survival package). To quantify prognostic risk, univariable and multivariable Cox proportional hazards regression models were constructed to calculate hazard ratios (HR) and 95% confidence intervals (CI). For multivariable analyses, models were adjusted for standard clinicopathological covariates tailored to the specific clinical cohort (detailed in the respective figure legends). A rule of 10 (± 5) events per covariate was followed to prevent model overfitting. Continuous variables were appropriately scaled prior to analysis; the continuous stromal signature and ODX scores were standardised via Z-score scaling, whereas age and tumour size were scaled to 10-year and 1-cm increments, respectively. Finally, in specific subgroups where perfect stratification occurred (zero events in the low-expression group), categorical Cox regression was precluded.

To evaluate the discriminatory accuracy of the continuous biomarkers, time-dependent ROC analysis with inverse probability of censoring weighting (IPCW) was performed using the timeROC package to generate area under the curve (AUC) metrics. Statistical significance was defined as a two-sided *P*-value < 0.05.

## Data and Code Availability

The spatial transcriptomic (GeoMx) data generated in this study, including raw DCC files and processed, normalised count matrices, have been deposited in the Gene Expression Omnibus (GEO) database under accession code GSE336314.

The bulk transcriptomic data and clinical metadata for the METABRIC validation cohort are publicly available through cBioPortal.

Any additional data supporting the findings of this study are available from the corresponding author upon reasonable request.

All custom R scripts and computational notebooks used for data processing, statistical analysis, survival modelling, and figure generation are contained in a private GitHub repository at github.com/ajordan003, available upon request.

