## Supplementary material for "Mural β3-integrin signposts Stroma6, a gene signature predicting early relapse in low-risk and chemoresistant breast cancer": Stroma6_Supplementary_BioRxiv_10Sep2026.pdf

**Extended Data Table 1.** Clinicopathological characteristics of the ODX NUH discovery, METABRIC, and NACT NUH validation cohorts

| **Characteristic** | **ODX NUH** ***n* = 302** | **METABRIC** ***n* = 785** | **NACT NUH** ***n* = 182** |
|---|---|---|---|
| ER Status | | | |
| Negative | 0 (0%) | 0 (0%) | 76 (42%) |
| Positive | 302 (100%) | 785 (100%) | 106 (58%) |
| HER2 Status | | | |
| Negative | 302 (100%) | 785 (100%) | 154 (85%) |
| Positive | 0 (0%) | 0 (0%) | 28 (15%) |
| Oncotype Risk Class | | | |
| High | 42 (14%) | 162 (21%) | - |
| Intermediate | 97 (32%) | 441 (56%) | - |
| Low | 163 (54%) | 182 (23%) | - |
| Relapse Status | | | |
| No Relapse | 274 (91%) | 532 (68%) | 90 (49%) |
| Relapsed | 28 (9.3%) | 253 (32%) | 92 (51%) |
| Follow-up Time (Months) | 48 (40 - 58) | 109 (64 - 164) | 63 (26 - 112) |
| Age at Diagnosis (Years) | 60 (50 - 68) | 65 (57 - 72) | 52 (46 - 61) |
| Unknown | 135 | 0 | 0 |
| Tumour Size (mm) | 21 (14 - 26) | 22 (18 - 30) | 22 (14 - 35) |
| Unknown | 137 | 6 | 0 |
| Tumour Grade | | | |
| Grade 1 | 15 (9.1%) | 88 (12%) | 3 (1.7%) |
| Grade 2 | 102 (62%) | 369 (49%) | 112 (62%) |
| Grade 3 | 48 (29%) | 297 (39%) | 65 (36%) |
| Unknown | 137 | 31 | 2 |
| Tumour Stage | | | |
| Stage 1 | 92 (56%) | 167 (29%) | 1 (0.5%) |
| Stage 2 | 71 (43%) | 384 (67%) | 35 (19%) |
| Stage 3 | 2 (1.2%) | 24 (4.2%) | 145 (80%) |
| Stage 4 | 0 (0%) | 0 (0%) | 1 (0.5%) |
| Unknown | 137 | 210 | 0 |
| Lymph Node Status | | | |
| Negative | 205 (68%) | 391 (50%) | 47 (26%) |
| Positive | 96 (32%) | 394 (50%) | 135 (74%) |
| Unknown | 1 | 0 | 0 |
| Neoadjuvant Chemotherapy | | | |
| Not Received | 302 (100%) | 785 (100%) | 0 (0%) |
| Received | 0 (0%) | 0 (0%) | 182 (100%) |
| Adjuvant Chemotherapy | | | |
| Not Received | 232 (77%) | 716 (91%) | - |
| Received | 70 (23%) | 69 (8.8%) | - |
| Unknown | 0 | 0 | - |
| Endocrine Therapy | | | |
| Not Received | 1 (0.3%) | 0 (0%) | 76 (42%) |
| Received | 296 (100%) | 785 (100%) | 104 (58%) |
| Unknown | 5 | 0 | 2 |

Data are presented as median (interquartile range) or *n* (%). Abbreviations: ODX, Oncotype DX; NUH, Nottingham University Hospital; NACT, neoadjuvant chemotherapy; ER, oestrogen receptor; HER2, human epidermal growth factor receptor 2; NA, not applicable; *n*, number of patients. 'Unknown' denotes missing retrospective clinical data for the specified variable.

**Extended Data Table 2.** Validation of the HALO AI αSMA⁺ BV classifier

| TMA Number | Batch | TMA Core | BVs Evaluated | Precision | Recall | F1-Score |
|---|---|---|---|---|---|---|
| TMA 1 | 1 | B8 | 25 | 1.00 | 0.87 | 0.93 |
| TMA 2 | 1 | H7 | 21 | 1.00 | 0.74 | 0.85 |
| TMA 1 | 2 | I6 | 22 | 1.00 | 0.92 | 0.96 |
| TMA 11 | 2 | J6 | 32 | 1.00 | 0.84 | 0.91 |
| Overall Performance | - | - | - | 1.00 | 0.84 | 0.91 |

Abbreviations: TMA, tissue microarray; BVs, blood vessels.

**Extended Data Table 3.** Prognostic performance of mural β3-integrin across clinical subgroups in the ODX NUH discovery cohort

| Subgroup | Time | *n* | Events | Log-rank *P* | Univariable HR (95% CI) | Cox *P* | AUC |
|---|---|---|---|---|---|---|---|
| ODX Low-Risk | 3-Year | 163 | 8 | < 0.001 | 15.80 (1.94-128.43) | 0.010 | 0.75 |
| ODX Intermediate-Risk | 3-Year | 97 | 3 | - | - | - | - |
| ODX High-Risk | 3-Year | 42 | 9 | 0.131 | 0.36 (0.09-1.44) | 0.149 | 0.43 |
| No Chemotherapy | 3-Year | 232 | 10 | 0.002 | 7.59 (1.61-35.75) | 0.010 | 0.70 |
| Chemotherapy | 3-Year | 70 | 10 | 0.163 | 2.95 (0.62-13.91) | 0.172 | 0.48 |

Statistics for the ODX intermediate-risk subgroup not calculated due to event sparsity. Abbreviations: ODX, Oncotype DX; NUH, Nottingham University Hospital; HR, hazard ratio; AUC, area under the curve; *n*, number of patients.

**Extended Data Table 4.** Prognostic evaluation of stromal (αSMA⁺) compartment-derived gene signatures (top 5-20) in the ODX NUH discovery cohort (low risk group) at 3 years

| Stromal Signature Genes | AIC | C-Index | 3-Year AUC | HR (Cont.) | *P* Value (Cont.) | HR (Cat.) | 95% CI | *P* Value (Cat.) | Log-Rank *P* Value |
|---|---|---|---|---|---|---|---|---|---|
| Top 5 | 74.77 | 0.76 | 0.76 | 3.18 | 0.027 | NE | NE | NE | 0.002 |
| Top 6 | 74.36 | 0.76 | 0.76 | 3.53 | 0.023 | NE | NE | NE | 0.002 |
| Top 7 | 75.92 | 0.73 | 0.73 | 2.74 | 0.043 | 8.80 | 1.08-71.50 | 0.042 | 0.014 |
| Top 8 | 75.68 | 0.73 | 0.74 | 2.97 | 0.039 | 8.04 | 0.99-65.39 | 0.051 | 0.020 |
| Top 9 | 75.53 | 0.75 | 0.75 | 3.33 | 0.036 | NE | NE | NE | 0.002 |
| Top 10 | 75.88 | 0.74 | 0.74 | 3.20 | 0.042 | NE | NE | NE | 0.001 |
| Top 11 | 75.92 | 0.74 | 0.74 | 3.22 | 0.043 | NE | NE | NE | 0.001 |
| Top 12 | 75.73 | 0.74 | 0.74 | 3.47 | 0.039 | NE | NE | NE | 0.002 |
| Top 13 | 75.63 | 0.75 | 0.75 | 3.80 | 0.037 | NE | NE | NE | 0.001 |
| Top 14 | 76.57 | 0.73 | 0.73 | 3.34 | 0.058 | NE | NE | NE | 0.002 |
| Top 15 | 76.59 | 0.73 | 0.73 | 3.33 | 0.060 | 8.29 | 1.02-67.36 | 0.048 | 0.018 |
| Top 16 | 76.88 | 0.72 | 0.72 | 3.17 | 0.069 | 8.54 | 1.05-69.40 | 0.045 | 0.016 |
| Top 17 | 76.90 | 0.71 | 0.71 | 3.27 | 0.069 | 8.29 | 1.02-67.36 | 0.048 | 0.018 |
| Top 18 | 77.41 | 0.70 | 0.70 | 2.90 | 0.090 | 8.29 | 1.02-67.36 | 0.048 | 0.018 |
| Top 19 | 77.29 | 0.71 | 0.71 | 3.08 | 0.085 | 7.81 | 0.96-63.48 | 0.055 | 0.023 |
| Top 20 | 77.15 | 0.71 | 0.71 | 3.24 | 0.079 | NE | NE | NE | 0.002 |

Stromal (αSMA⁺) signatures were evaluated as both continuous (cont.) variables (signature score) and categorical (cat.) variables (median stratification) in the ODX NUH discovery cohort low-risk group (*n* = 142). Where not estimable (NE is shown), standard categorical Cox proportional hazards models failed to converge due to zero relapse events in the low-expression group. Abbreviations: ODX, Oncotype DX; NUH, Nottingham University Hospital; AIC, Akaike information criterion; C-Index, Harrell's concordance index; AUC, area under the curve; HR, hazard ratio; CI, confidence interval; Cont., continuous; Cat., categorical.

**Extended Data Table 5.** Prognostic evaluation of tumour (PanCK⁺) compartment-derived gene signatures (top 5-20) in the ODX NUH discovery cohort (low risk group) at 3 years

| Tumour Signature Genes | AIC | C-Index | 3-Year AUC | HR (Cont.) | *P* Value (Cont.) | HR (Cat.) | 95% CI | *P* Value (Cat.) | Log-Rank *P* Value |
|---|---|---|---|---|---|---|---|---|---|
| Top 5 | 70.25 | 0.62 | 0.62 | 1.39 | 0.226 | 1.32 | 0.30-5.89 | 0.718 | 0.717 |
| Top 6 | 70.12 | 0.62 | 0.62 | 1.50 | 0.207 | 1.32 | 0.30-5.89 | 0.718 | 0.717 |
| Top 7 | 70.20 | 0.60 | 0.60 | 1.48 | 0.219 | 1.36 | 0.30-6.06 | 0.691 | 0.690 |
| Top 8 | 70.32 | 0.60 | 0.60 | 1.53 | 0.241 | 1.39 | 0.31-6.23 | 0.664 | 0.663 |
| Top 9 | 70.27 | 0.61 | 0.61 | 1.61 | 0.234 | 1.43 | 0.32-6.40 | 0.638 | 0.636 |
| Top 10 | 69.50 | 0.65 | 0.64 | 1.84 | 0.137 | 1.28 | 0.29-5.73 | 0.745 | 0.744 |
| Top 11 | 69.53 | 0.64 | 0.64 | 1.93 | 0.141 | 1.25 | 0.28-5.57 | 0.773 | 0.772 |
| Top 12 | 69.57 | 0.64 | 0.64 | 2.01 | 0.145 | 1.21 | 0.27-5.42 | 0.800 | 0.800 |
| Top 13 | 69.41 | 0.63 | 0.64 | 2.09 | 0.133 | 1.28 | 0.29-5.73 | 0.745 | 0.744 |
| Top 14 | 69.33 | 0.64 | 0.64 | 2.12 | 0.127 | 1.39 | 0.31-6.23 | 0.664 | 0.663 |
| Top 15 | 69.26 | 0.63 | 0.63 | 2.21 | 0.120 | 1.36 | 0.30-6.06 | 0.691 | 0.690 |
| Top 16 | 69.43 | 0.63 | 0.63 | 2.22 | 0.134 | 1.39 | 0.31-6.23 | 0.664 | 0.663 |
| Top 17 | 69.69 | 0.62 | 0.62 | 2.11 | 0.158 | 1.39 | 0.31-6.23 | 0.664 | 0.663 |
| Top 18 | 69.68 | 0.61 | 0.61 | 2.18 | 0.156 | 1.43 | 0.32-6.40 | 0.638 | 0.636 |
| Top 19 | 69.80 | 0.61 | 0.61 | 2.19 | 0.170 | 1.36 | 0.30-6.06 | 0.691 | 0.690 |
| Top 20 | 69.56 | 0.62 | 0.62 | 2.38 | 0.145 | 1.43 | 0.32-6.40 | 0.638 | 0.636 |

Tumour (PanCK⁺) signatures were evaluated as both continuous (cont.) variables (signature score) and categorical (cat.) variables (median stratification) in the ODX NUH discovery cohort low-risk group ($n$ = 151). Abbreviations: ODX, Oncotype DX; NUH, Nottingham University Hospital; AIC, Akaike information criterion; C-Index, Harrell's concordance index; AUC, area under the curve; HR, hazard ratio; CI, confidence interval; Cont., continuous; Cat., categorical.

**Extended Data Table 6.** Prognostic evaluation of immune (CD45⁺) compartment-derived gene signatures (top 5-20) in the ODX NUH discovery cohort (low risk group) at 3 years

| Immune Signature Genes | AIC | C-Index | 3-Year AUC | HR (Cont.) | *P* Value (Cont.) | HR (Cat.) | 95% CI | *P* Value (Cat.) | Log-Rank *P* Value |
|---|---|---|---|---|---|---|---|---|---|
| Top 5 | 61.45 | 0.67 | 0.68 | 5.48 | 0.028 | 1.86 | 0.42-8.30 | 0.418 | 0.411 |
| Top 6 | 61.45 | 0.67 | 0.69 | 6.62 | 0.031 | 1.70 | 0.38-7.61 | 0.485 | 0.480 |
| Top 7 | 63.33 | 0.61 | 0.62 | 5.14 | 0.114 | 1.49 | 0.33-6.67 | 0.601 | 0.598 |
| Top 8 | 63.53 | 0.62 | 0.63 | 5.37 | 0.133 | 0.85 | 0.19-3.82 | 0.836 | 0.836 |
| Top 9 | 63.72 | 0.62 | 0.64 | 5.26 | 0.147 | 1.63 | 0.37-7.30 | 0.521 | 0.517 |
| Top 10 | 63.94 | 0.61 | 0.63 | 4.68 | 0.164 | 3.14 | 0.61-16.19 | 0.172 | 0.149 |
| Top 11 | 63.02 | 0.66 | 0.68 | 7.30 | 0.091 | 2.77 | 0.54-14.26 | 0.224 | 0.205 |
| Top 12 | 62.98 | 0.66 | 0.68 | 8.08 | 0.089 | 2.88 | 0.56-14.87 | 0.206 | 0.185 |
| Top 13 | 63.39 | 0.62 | 0.63 | 6.45 | 0.110 | 1.57 | 0.35-6.99 | 0.558 | 0.554 |
| Top 14 | 63.06 | 0.61 | 0.63 | 6.39 | 0.084 | 2.71 | 0.53-14.00 | 0.233 | 0.214 |
| Top 15 | 63.57 | 0.60 | 0.61 | 4.83 | 0.119 | 1.37 | 0.31-6.12 | 0.680 | 0.679 |
| Top 16 | 64.33 | 0.58 | 0.60 | 3.32 | 0.215 | 1.26 | 0.28-5.63 | 0.762 | 0.762 |
| Top 17 | 64.17 | 0.60 | 0.61 | 3.46 | 0.186 | 1.26 | 0.28-5.63 | 0.762 | 0.762 |
| Top 18 | 63.90 | 0.60 | 0.61 | 4.09 | 0.154 | 1.37 | 0.31-6.12 | 0.680 | 0.679 |
| Top 19 | 63.55 | 0.60 | 0.61 | 4.94 | 0.121 | 1.26 | 0.28-5.63 | 0.762 | 0.762 |
| Top 20 | 63.99 | 0.58 | 0.59 | 4.03 | 0.170 | 1.31 | 0.29-5.87 | 0.721 | 0.720 |

Immune (CD45⁺) signatures were evaluated as both continuous (cont.) variables (signature score) and categorical (cat.) variables (median stratification) in the ODX discovery cohort low-risk group ($n$ = 99). Abbreviations: ODX, Oncotype DX; NUH, Nottingham University Hospital; AIC, Akaike information criterion; C-Index, Harrell's concordance index; AUC, area under the curve; HR, hazard ratio; CI, confidence interval; Cont., continuous; Cat., categorical.

**Extended Data Table 7.** Prognostic performance of Stroma6 across clinical subgroups in the ODX NUH discovery cohort

| Subgroup | Time | *n* | Events | Log-rank *P* | Univariable HR (95% CI) | Cox *P* | AUC |
|---|---|---|---|---|---|---|---|
| ODX Low-Risk | 3-Year | 142 | 8 | 0.002 | NE | NE | 0.76 |
| ODX Intermediate-Risk | 3-Year | 90 | 3 | 0.550 | 2.05 (0.19-22.59) | 0.558 | 0.65 |
| ODX High-Risk | 3-Year | 38 | 7 | 0.085 | 0.26 (0.05-1.36) | 0.111 | 0.31 |
| No Chemotherapy | 3-Year | 206 | 10 | 0.006 | 10.38 (1.31-81.90) | 0.026 | 0.73 |
| Chemotherapy | 3-Year | 64 | 8 | 0.238 | 0.43 (0.10-1.82) | 0.253 | 0.37 |

Where not estimable (NE is shown), standard categorical Cox proportional hazards models failed to converge due to zero relapse events in the low-expression group. Abbreviations: ODX, Oncotype DX; NUH, Nottingham University Hospital; HR, hazard ratio; AUC, area under the curve; *n*, number of patients.

**Extended Data Table 8.** Prognostic performance of Stroma6 across clinical subgroups in the curated METABRIC cohort

| Subgroup | Time | *n* | Events | Log-rank *P* | Univariable HR (95% CI) | Cox *P* | AUC |
|---|---|---|---|---|---|---|---|
| ODX Low-Risk | 3-Year | 182 | 9 | < 0.001 | 9.91 (2.06-47.70) | 0.004 | 0.71 |
| ODX Intermediate-Risk | 3-Year | 441 | 32 | 0.479 | 0.78 (0.39-1.56) | 0.480 | 0.50 |
| ODX High-Risk | 3-Year | 162 | 32 | 0.068 | 2.02 (0.93-4.36) | 0.074 | 0.61 |
| No Chemotherapy | 3-Year | 716 | 68 | 0.006 | 1.97 (1.20-3.22) | 0.007 | 0.62 |
| Chemotherapy | 3-Year | 69 | 5 | 0.491 | 0.47 (0.05-4.22) | 0.501 | 0.39 |

Abbreviations: ODX, Oncotype DX; HR, hazard ratio; AUC, area under the curve; *n*, number of patients.

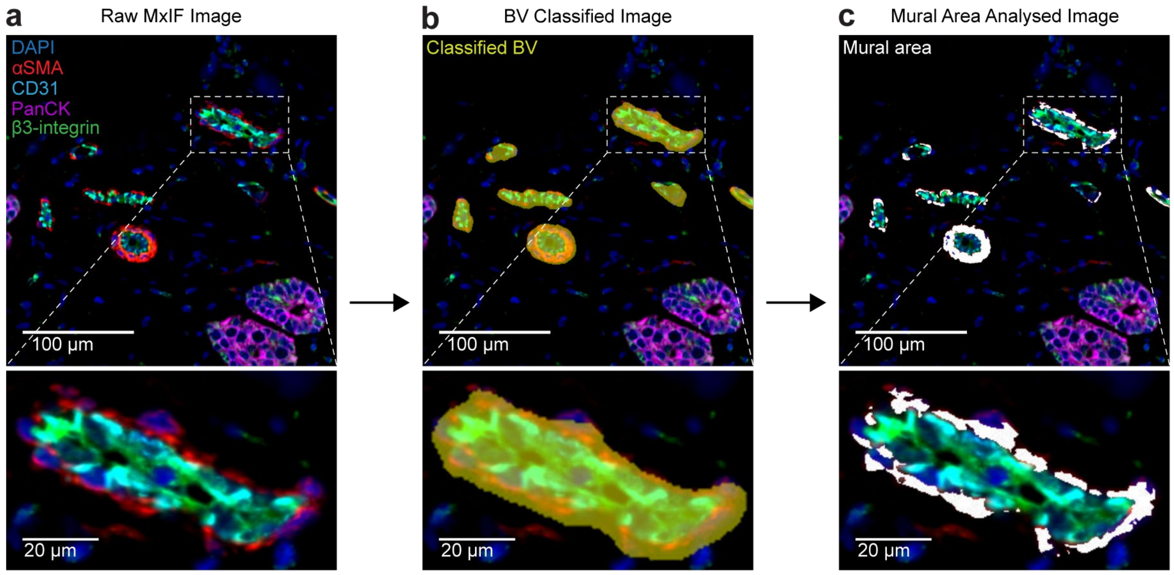


**Extended Data Figure 1. HALO AI quantification of mural β3-integrin**

Image analysis workflow in the discovery cohort. **a**, Representative Cell DIVE multiplex immunofluorescence (MxIF) images display DAPI (blue), αSMA (red), CD31 (aqua), PanCK (magenta), and β3-integrin. **b**, A HALO AI classifier segmented $\alpha SMA^+$ tumour blood vessels (yellow overlay). **c**, β3-integrin intensity was subsequently quantified specifically within the $\alpha SMA^+/CD31^-$ mural compartment (white mask). Scale bars = 100 μm; insets = 20 μm.

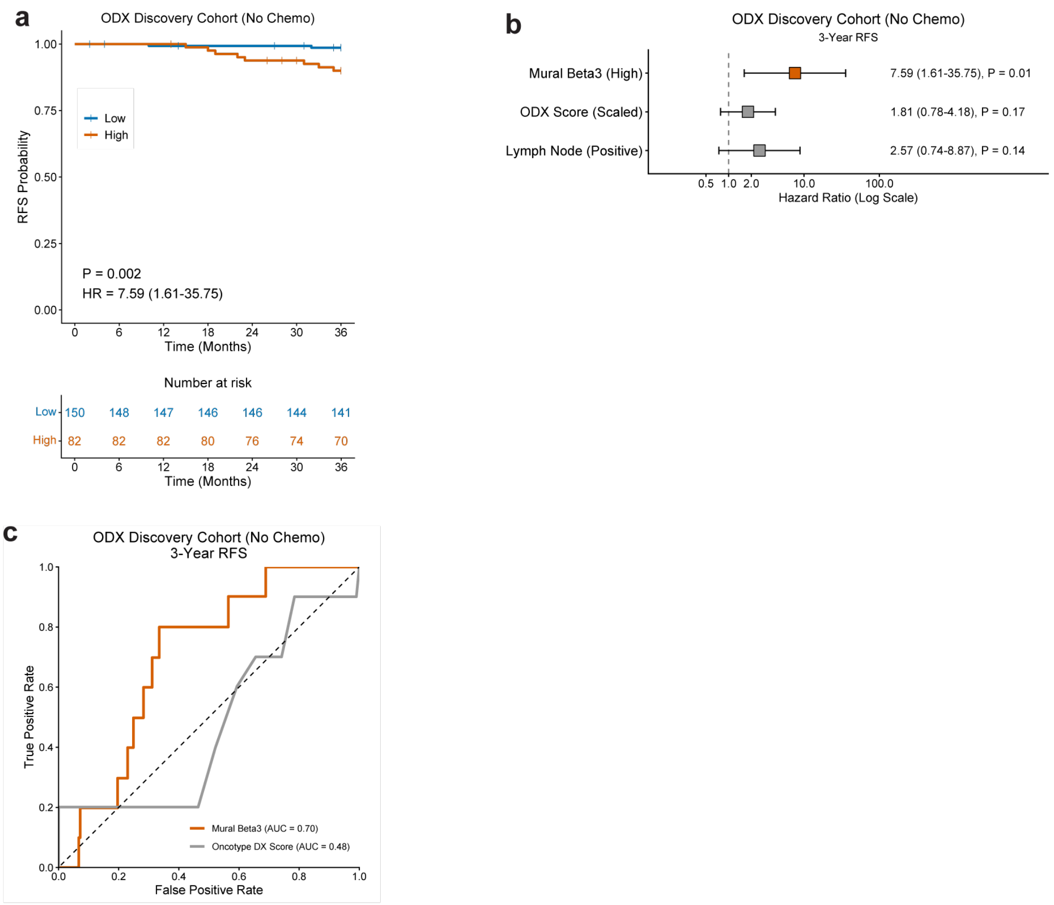


**Extended Data Figure 2. High mural β3-integrin protein expression predicts early relapse in chemotherapy-naive breast cancer**

**a**, Kaplan–Meier (KM) 3-year relapse-free survival (RFS) comparing high (orange) and low (blue) mural β3-integrin in the Oncotype DX (ODX) Nottingham University Hospital (NUH) discovery cohort no adjuvant chemotherapy group (*n* = 232). *P* values by two-sided log-rank test. **b**, Univariable Cox regression for 3-year RFS in the ODX NUH no adjuvant chemotherapy cohort (*n* = 232). HR shown for high mural β3-integrin (reference: low), positive lymph node (LN) invasion (reference: negative), and ODX recurrence score (RS), where HR reflects a 1-standard deviation (SD, 11.1 points) increase. Error bars represent 95% confidence intervals. **c**, Time-dependent receiver operating characteristic (ROC) curves comparing continuous mural β3-integrin (orange) and ODX RS (grey) for 3-year relapse in the ODX NUH no adjuvant chemotherapy cohort (*n* = 232). AUC, area under the curve. Patients were stratified into high and low mural β3-integrin groups using cut-off points derived by maximally selected rank statistics.

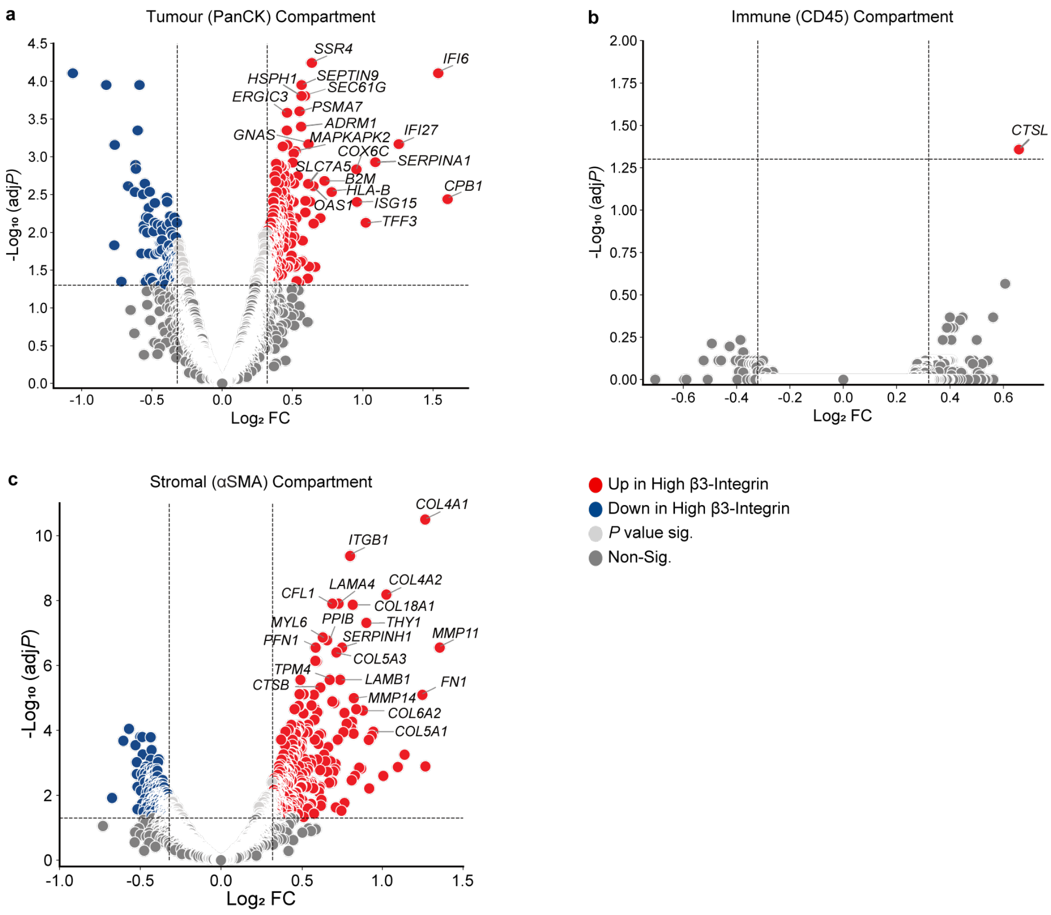


**Extended Data Figure 3. Compartment-specific differential gene expression by mural β3-integrin status**

**a–c**, Volcano plots of differential gene expression (two-sided moderated t-test, Benjamini–Hochberg correction) between mural β3-integrin-high and -low tumours in the Oncotype DX (ODX) Nottingham University Hospital (NUH) discovery cohort low-risk group. Analyses are shown for the (**a**) tumour (PanCK$^+$; $n$ = 144), (**b**) immune (CD45$^+$; $n$ = 94), and (**c**) stromal (αSMA$^+$; $n$ = 137) compartments. Coloured points denote significant up- (red) or down-regulation (blue) when $Log_2$ fold change (FC) ≥ 0.32 and adjusted (adj) $P$ value ≤ 0.05.

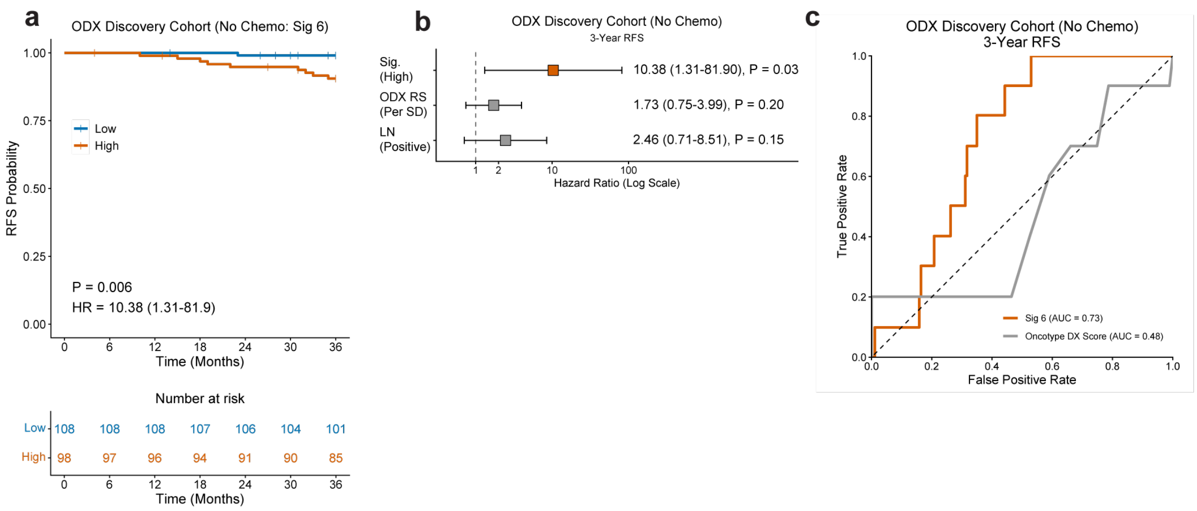


**Extended Data Figure 4. Stroma6 high signature predicts early relapse in chemotherapy-naive breast cancer**

**a**, Kaplan–Meier (KM) 3-year relapse-free survival (RFS) comparing high (orange) and low (blue) Stroma6 in the Oncotype DX (ODX) Nottingham University Hospital (NUH) discovery cohort no adjuvant chemotherapy group (*n* = 206). P values by two-sided log-rank test. **b**, Univariable Cox regression for 3-year RFS in the ODX NUH no adjuvant chemotherapy cohort (*n* = 206). HR shown for high Stroma6 (reference: low), positive lymph node (LN) invasion (reference: negative), and ODX recurrence score (RS), where HR reflects a 1-standard deviation (SD, 11.1 points) increase. Error bars represent 95% confidence intervals. **c**, Time-dependent receiver operating characteristic (ROC) curves comparing continuous Stroma6 score (orange) and ODX RS (grey) for 3-year relapse in the ODX NUH no adjuvant chemotherapy cohort (*n* = 206). Patients were stratified into high and low Stroma6 groups using the global median Stroma6 score of the full, unstratified dataset. AUC, area under the curve.

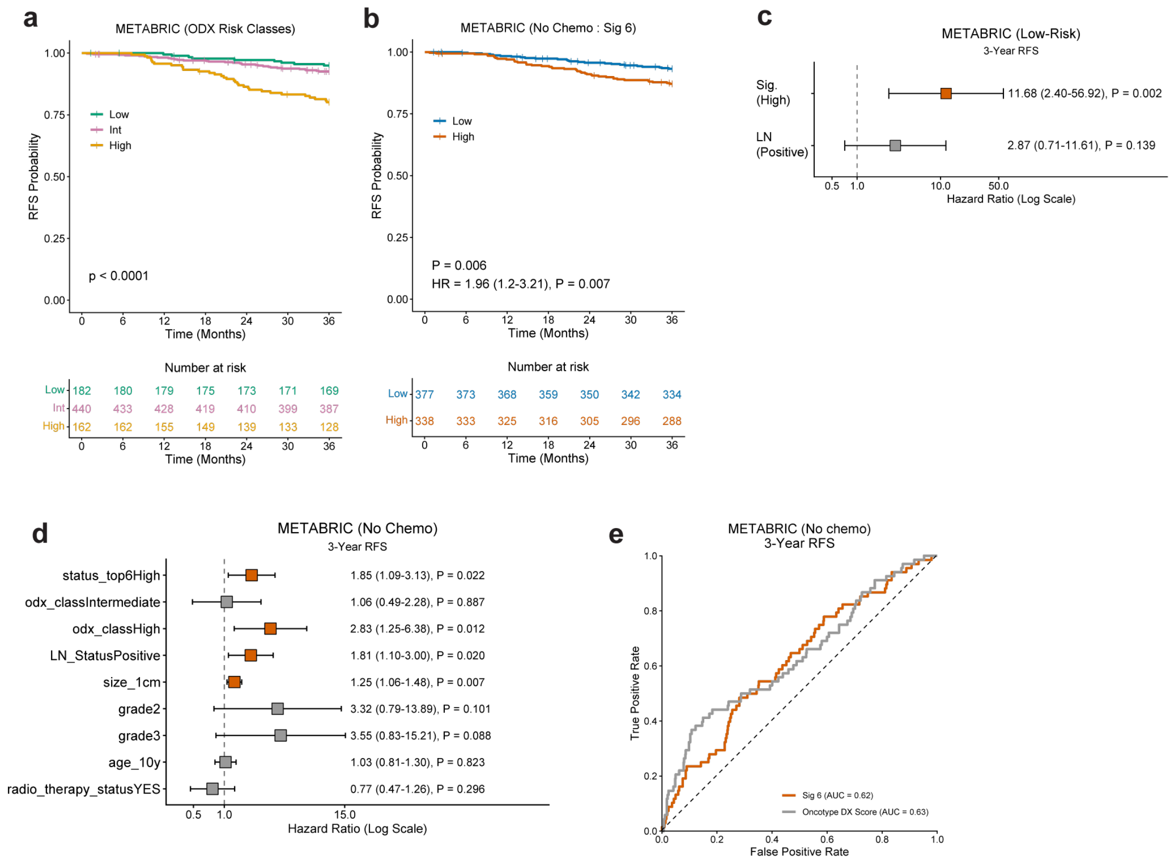


**Extended Data Figure 5. A stromal gene signature predicts early relapse in Oncotype DX low-risk and chemotherapy-naive patients within the METABRIC cohort.**

**a,** Kaplan–Meier (KM) 3-year relapse-free survival (RFS) comparing in silico Oncotype DX (ODX) risk classes (low = green, intermediate = pink, high = yellow) in the $ER^+/HER2^-$ METABRIC external cohort (*n* = 784). **b**, KM 3-year RFS comparing high (orange) and low (blue) Stroma6 in the no adjuvant chemotherapy group (*n* = 715). **c–d**, Multivariable Cox regression for (**c**) 3-year RFS in the ODX low-risk group (*n* = 182), and (**d**) 3-year no adjuvant chemotherapy group (*n* = 682). HR shown for high Stroma6 (reference: low), positive lymph node (LN) invasion (reference: negative), tumour size (per 1 cm), ODX risk class (intermediate, high; reference: low), tumour grade (grade 2, grade 3; reference: grade 1), age (per 10 years), and adjuvant radiotherapy (reference: not received). Error bars represent 95% confidence intervals. **e**, Time-dependent receiver operating characteristic (ROC) curves comparing continuous Stroma6 score (orange) and ODX recurrence score (RS; grey) for 3-year in the no adjuvant chemotherapy cohort (*n* = 715). AUC, area under the curve. For **a–b**, *P* value by two-sided log-rank test. Patients were stratified into high and low Stroma6 groups using the global median Stroma6 score of the full, unstratified dataset

**a**

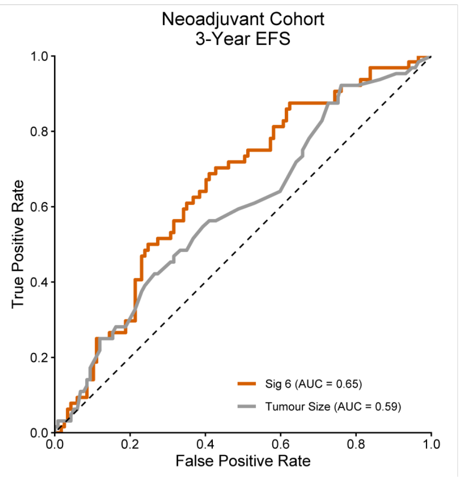


**Extended Data Figure 6. Stroma6 gene signature predicts early relapse in chemoresistant breast cancer**

**a,** Time-dependent receiver operating characteristic (ROC) curves comparing continuous Stroma6 score (orange) and tumour size (grey) for 3-year EFS in the neoadjuvant chemotherapy (NACT) Nottingham University Hospital (NUH) cohort (*n* = 182). AUC, area under the curve.
